# CCP1, a tubulin deglutamylase, increases survival of rodent spinal cord neurons following glutamate-induced excitotoxicity

**DOI:** 10.1101/2020.09.16.295279

**Authors:** Yasmin H. Ramadan, Amanda Gu, Nicole Ross, Sara A. McEwan, Maureen M. Barr, Bonnie L. Firestein, Robert O’Hagan

## Abstract

Microtubules (MTs) are cytoskeletal elements that provide structural support, establish morphology, and act as roadways for intracellular transport in cells. Neurons extend and must maintain long axons and dendrites to transmit information through the nervous system. Therefore, in neurons, the ability to independently regulate cytoskeletal stability and MT-based transport in different cellular compartments is essential. Post-translational modification of MTs is one mechanism by which neurons can regulate the cytoskeleton.

The carboxypeptidase CCP1 negatively regulates post-translational glutamylation of MTs. We previously demonstrated that the CCP1 homolog in *C. elegans* is important for maintenance of cilia. In mammals, loss of CCP1, and the resulting hyperglutamylation of MTs, causes neurodegeneration. It has long been known that CCP1 expression is activated by neuronal injury; however, whether CCP1 plays a neuroprotective role after injury is unknown. Furthermore, it not yet clear whether CCP1 acts on ciliary MTs in spinal cord neurons.

Using an *in vitro* model of excitotoxic neuronal injury coupled with shRNA-mediated knockdown of CCP1, we demonstrate that CCP1 protects neurons from excitotoxic death. Unexpectedly, excitotoxic injury reduced CCP1 expression in our system, and knockdown of CCP1 did not result in loss or shortening of cilia in cultured spinal cord neurons. Our results suggest that CCP1 acts on axonal and dendritic MTs to promote cytoskeletal rearrangements that support neuroregeneration and that enzymes responsible for glutamylation of MTs might be therapeutically targeted to prevent excitotoxic death after spinal cord injuries.

## Introduction

The development, function, and survival of neurons rely heavily on the function of the MT cytoskeleton [1, 2]. MTs are hollow cylinders formed by polymerization of α and β tubulin subunits [3]. Tubulins are highly conserved, differing little in sequence and structure [4]. Yet, MTs in different neuronal compartments, such as axons, dendrites, or growth cones, display differences in function and dynamics [2]. The Tubulin Code model proposes that in addition to heterogeneity of tubulin isotype composition, post-translational modification of tubulins can specialize the stability, form, and function of MTs [4, 5]. Tubulin Code modifications are proposed to endow specific MTs with particular properties to play essential roles in the function of axons and dendrites, as well as directional trafficking, plasticity, and survival [4, 5]. Glutamylation, one component of the Tubulin Code, consists of side-chains of the amino acid glutamate that are post-translationally added to the carboxy terminal tails of tubulins when assembled into MTs [6, 7]. Glutamylation often decorates MTs in both neurons and cilia [4].

The primary cilium, a non-motile sensory organelle that protrudes from most non-dividing cells in the human body, is also a region of MT specialization [8]. Conserved over eukaryotic evolution from algae to vertebrates, the architecture of cilia consists of a MT cytoskeleton, called the axoneme, in which a ring of nine MT doublets extends along cilia immediately beneath the membrane [8]. Vertebrate cilia have been studied in processes such as kidney function, olfaction, vision, and development of left-right asymmetry [9]. Cilia also play a role in nervous system development due to their function as an essential hub for signaling pathways [10, 11].

Glutamylation regulates the structure and function of cilia [12–14]. M14D carboxypeptidases, such as CCP1 [15], remove or reduce the length of glutamate side-chains [16]. When deglutamylase function is lost, hyperglutamylation affects ciliary motor transport and causes degeneration of some types of neuronal sensory cilia in *C. elegans* [13, 17]. In mice, loss of CCP1 leads to the degeneration of retinal photoreceptors and sperm defects [18]. These phenotypes are reminiscent of the symptoms of diseases caused by ciliary dysfunction, or “ciliopathies” [19].

Glutamylation also occurs on non-ciliary neuronal MTs [20]. Loss of CCP1 perturbs neuronal transport in mice and humans [18, 21, 22] and leads to infantile hereditary neurodegeneration and cerebellar atrophy in humans [23]. In mammals, expression of CCP1 deglutamylase is upregulated in response to transection or crush injury of the sciatic nerve, suggesting that its function may be required for neuroregeneration [24]. Loss of deglutamylase activity diminishes regrowth of laser-severed neurons in *C. elegans*, supporting a possible conserved role in neuroregeneration [25].

Questions about the function of CCP1 remain unanswered. Is CCP1 neuroprotective or does it play a pathological role after neuronal injury? Is CCP1 expression activated in injured neurons of the central nervous system (CNS), as it is in the sciatic nerve in the peripheral nervous system (PNS)? Does CCP1 function in cilia, somata, or neurites of injured spinal cord neurons?

Here, we tested if CCP1 activity affects neuronal cilia in mammals by analyzing cilia in embryonic rat spinal cord cultures using shRNA to knock down CCP1 expression. Immunofluorescence-based detection of the ciliary marker ARL13B suggests that a primary cilium protruded from the majority of neurons in our spinal cord cultures. Unexpectedly, CCP1 knockdown increased, rather than decreased, the percentage of ciliated neurons, suggesting that CCP1 may promote ciliation in cultured rat spinal cord neurons.

Using an *in vitro* model of the secondary phase of spinal cord injury [26], we also found that knockdown of CCP1 decreased the survival of spinal cord neurons subjected to excitotoxic glutamate treatment, suggesting that CCP1 is neuroprotective. In contrast to the reported upregulation of CCP1 in response to injury of the sciatic nerve, our analysis showed that glutamate-induced excitotoxic injury reduced CCP1 expression in spinal cord neurons. However, shRNA CCP1 knockdown in cells subjected to excitotoxic glutamate did not reduce CCP1 expression to lower levels than excitoxic glutamate alone. Therefore, we propose that the neuroprotective activity of CCP1 must occur before excitoxic injury, most likely by acting on MTs in non-ciliary compartments, such as axons, dendrites, or somata.

## Results

### Mammalian spinal cord neurons in culture are ciliated

Because glutamylation is implicated in degeneration or dysfunction of cilia and flagella [13, 17, 18, 27–31], we tested if CCP1 knockdown influences ciliation of spinal cord neurons. We first assessed if spinal cord neurons and glia from cultured rat embryonic spinal cord are ciliated. Cilia of spinal cord neurons have not been well-documented. To our knowledge, only the CSF-contacting neurons [32–35], neuronal precursors and ependymal cells that line the central canal [36], and motor neurons of the lumbar spinal cord [37] are known to be ciliated. To test the role of CCP1 in spinal cord neuron ciliation, we isolated and dissociated embryonic rat spinal cords **(Fig. 1A)** from pregnant mothers at gestational day 15 (E15) and grew mixed cultures of astrocytes, microglia, and neurons (**Fig. 1B**). Indirect immunofluorescence using an antibody specific to the ciliary membrane protein ARL13B [38] demonstrated that 57% of spinal cord neurons in our cultures were ciliated at DIV (day *in vitro*) 8 (**Fig. 2A, B**). Cholinergic motor neurons do not survive in embryonic spinal cord cultures under similar culture conditions [39]. Neurons in our cultures lacked immunostaining against ChAT (**Fig. S1A**), and therefore were most likely sensory neurons and interneurons, not motorneurons. Assuming that sensory and interneurons in our rat spinal cord cultures are similar to spinal cords neurons *in vivo,* we conclude that sensory neurons of the spinal cord are typically ciliated.

**Fig. 1.**
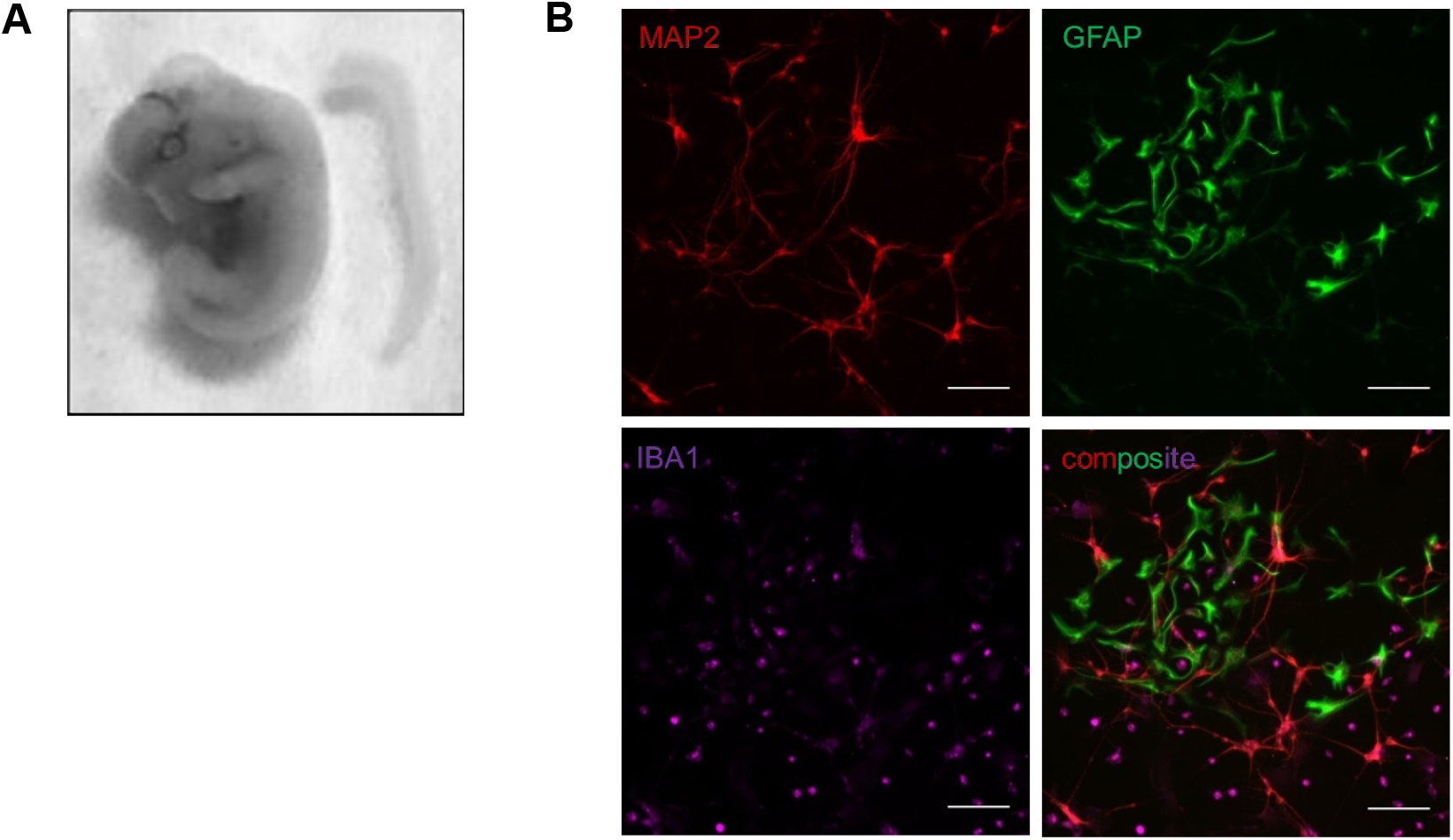
Embryonic spinal cord cultures contained a mixture of neurons, astrocytes, and microglia at DIV7. **A.** Image of a rat embryo collected at embryonic day 15 (left) and a dissected spinal cord at this developmental stage (right). **B.** Primary spinal cord cultures derived from embryos at gestational day 15 contained neurons (MAP2+), astrocytes (GFAP+), and microglia (IBA1+) at DIV7, as labelled by immunocytochemistry. Scale bars indicate 100μm.

**Fig. 2.**
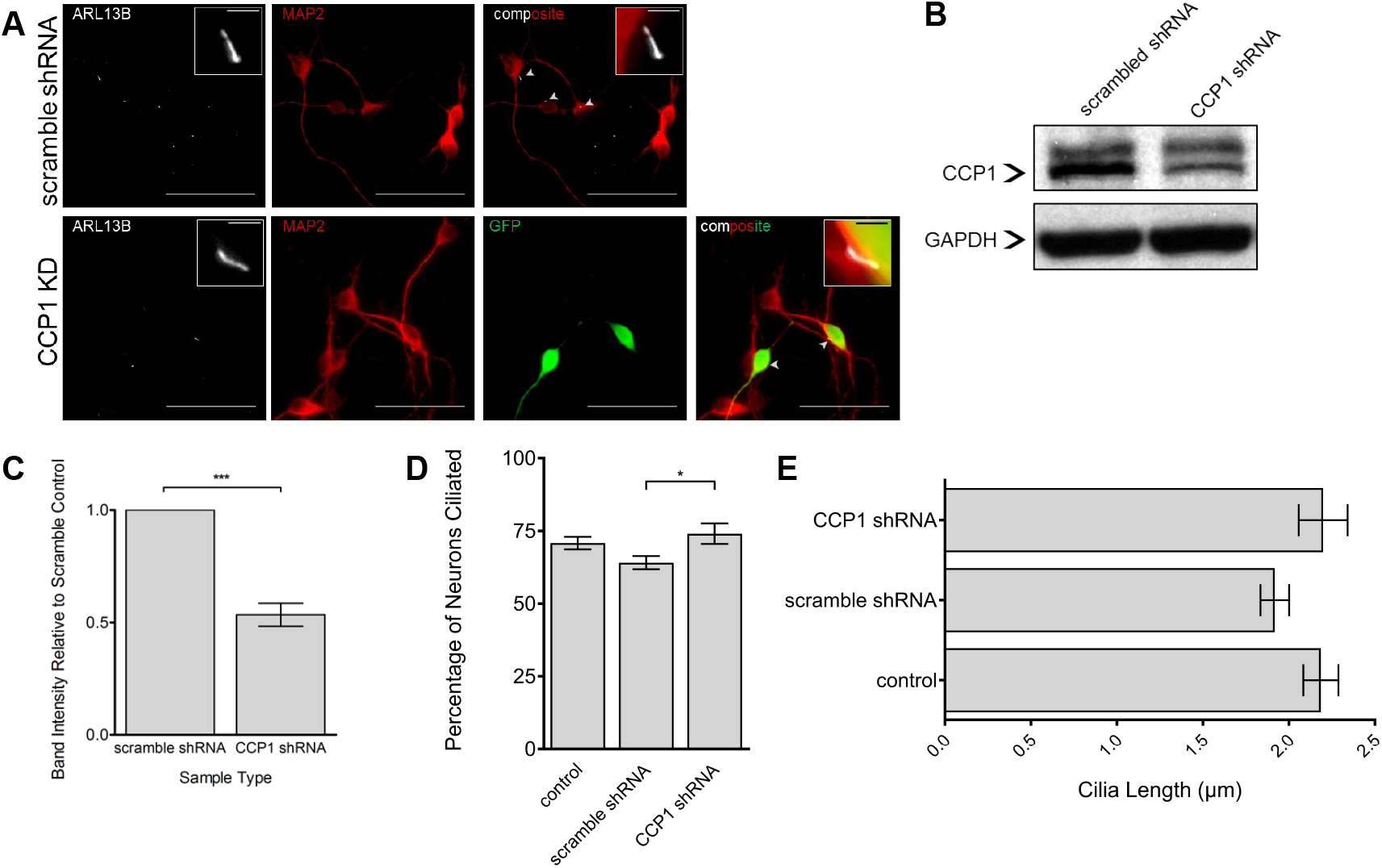
Knockdown of CCP1 increases incidence of ciliated spinal cord neurons. **A.** Primary spinal cord cultures immunostained for neuronal marker MAP2 and cilia marker ARL13B, showing ciliated neurons. shRNA knockdown (scramble or CCP1) is indicated. Scale bars indicate 50μm and 2.5μm (inset). Arrowheads point to cilia. See also **Fig. S1**. **B.** Western blot showing CCP1 and GAPDH (loading control) in primary spinal cord cultures transduced with scrambled shRNA or CCP1 shRNA. **C.** Quantification of CCP1 knockdown, error bars indicate SEM. \*\*\**p* < 0.001, determined by paired t-test. n=3. **D.** Quantification of percentage of neurons that were ciliated in control (no shRNA), scramble shRNA control, and CCP1 shRNA knockdown cultures, \*\**p* < 0.01 determined by Student’s t-test. **E.** Quantification of neuronal cilia length. Cilia length was not significantly different in control, scramble shRNA control, or CCP1 shRNA knockdown cultures (Student’s t-test).

### Knockdown of the deglutamylase CCP1 increases the presence of cilia on spinal cord neurons, but does not significantly affect ciliary length

We hypothesized that if CCP1 functions to maintain cilia in mammalian spinal cord neurons, as it does in *C. elegans*, then loss of CCP1 function would cause ciliary degeneration [13]. To test this, we prepared cultures of rat spinal cords and used a lentivirus containing an shRNA sequence targeting the 3’UTR of CCP1 to knock down CCP1 expression. shRNA knockdown of CCP1 resulted in a 50% decrease of CCP1 protein levels compared to cultures infected with a lentivirus containing a scrambled shRNA sequence (**Fig. 2B, C**). As expected, the lentivirus containing shRNA specific to CCP1 selectively infected more neurons than glia (non-neuronal cells) (**Fig. S1B, C**). Therefore, the shRNA-mediated decrease in CCP1 levels in these spinal cord cultures is most likely due to knockdown of CCP1 in neurons.

Next, we counted cilia present on neurons and measured their lengths. Surprisingly, we found that shRNA-mediated knockdown of CCP1 increased the frequency of ciliated neurons (**Fig. 2D**). However, knockdown of CCP1 did not affect ciliary length (**Fig. 2E**). Thus, our data suggests that reduction of CCP1 might not lead to degeneration of primary cilia in the murine spinal cord, as it does in *C. elegans* sensory neurons, at least on a time scale of several days.

### Knockdown of CCP1 decreases neuronal survival of excitotoxic challenge

Loss of CCP1 leads to severe neuronal degeneration in the murine central nervous system (CNS) [18]. Additionally, a recent study demonstrated that loss of CCP1 in humans results in degeneration of cerebellar neurons and spinal neurons [23]. Therefore, we sought to test how knockdown of CCP1 function affects the survival of spinal cord neurons.

We subjected our cultures to glutamate-induced excitotoxicity, which is characteristic of the secondary stage of spinal cord injury [26], by incubating the cultures with concentrations of glutamate ranging from 200μM to 1mM for 1 hour on DIV 7 (**Fig. 3A)**. After injury, cultures were allowed to recover for 24 hours before fixation on DIV 8 and subsequent immunostaining for MAP2. MAP2+ surviving neurons were counted, and as expected, glutamate exposure caused neuronal death in a dose-dependent manner (**Fig. 3B, C**). Approximately, 60% of neurons survived following exposure to 1mM glutamate for 1 hour.

**Fig. 3.**
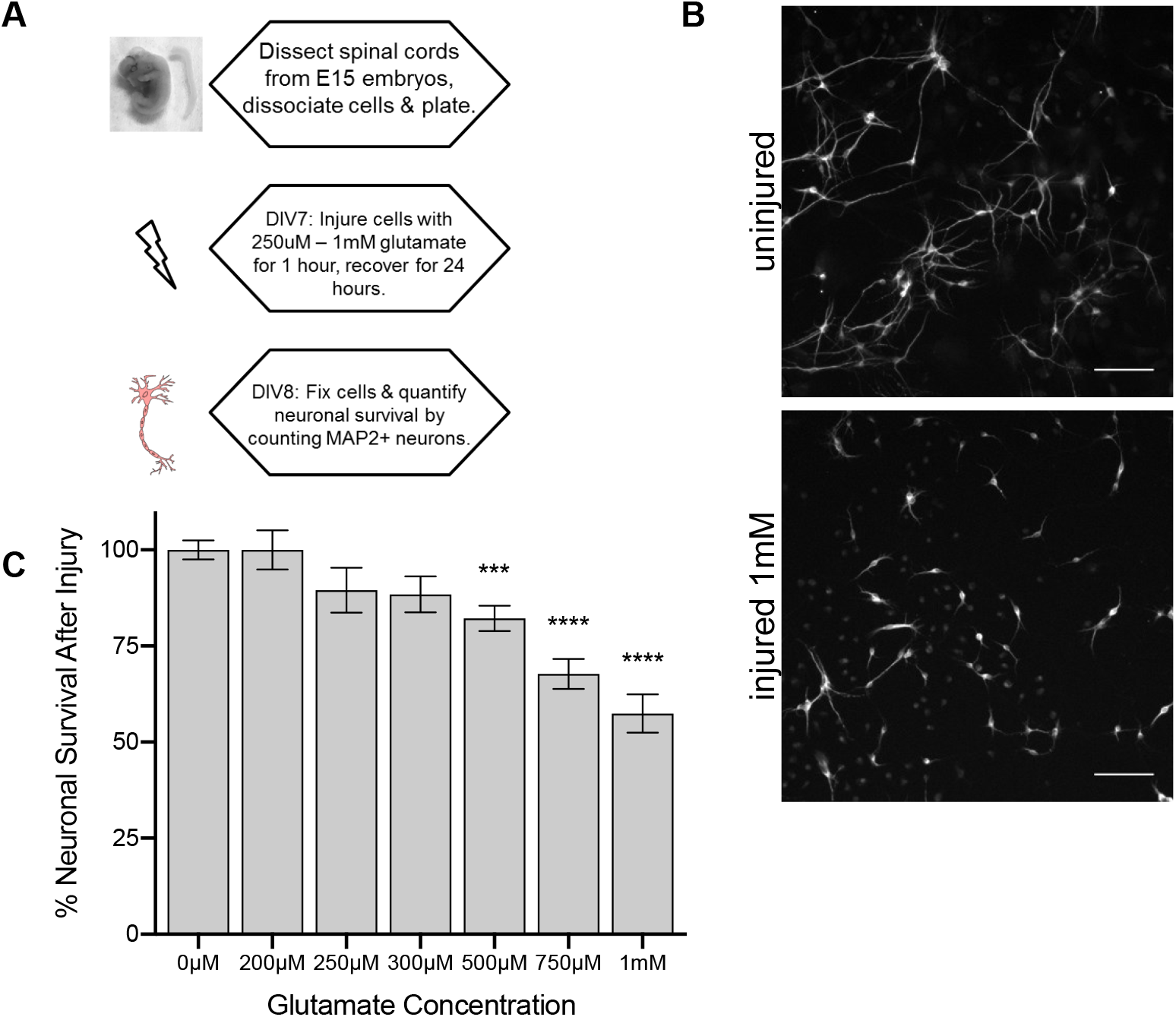
Neuronal death following glutamate-induced excitotoxicity is modeled *in vitro*. **A.** Schematic of treatment of spinal cord cultures with excitotoxic glutamate. Primary spinal cord cultures containing neurons and non-neuronal glial cells were treated with varying concentrations of glutamate for 1hour on DIV7 and allowed to recover for 24 hours before fixation to mimic glutamate-induced excitotoxicity following spinal cord injury. **B.** Control and injured spinal cord cultures were immunostained for MAP2 to identify neurons. Glutamate-induced excitotoxic injury led to loss of neurons and retracted neuronal processes compared with uninjured neurons. Scale bars indicate 100μm. **C.** Percent neuronal survival after excitotoxic injury decreased with increasing glutamate concentration. Error bars represent SEM. *** indicates *p*<0.001, **** indicates *p*<0.0001 versus 0μM glutamate by one-way ANOVA and Dunnett’s Multiple Comparison Test.

To determine whether CCP1 is required for neuronal survival following glutamate-induced excitotoxicity, cultures in which CCP1 levels were knocked down by lentiviral shRNA treatment were subjected to varying concentrations of glutamate. Neuronal survival following glutamate exposure was significantly reduced in cultures transduced with shRNA CCP1 knock down at all glutamate concentrations tested (**Fig. 4A, B**).

**Fig. 4.**
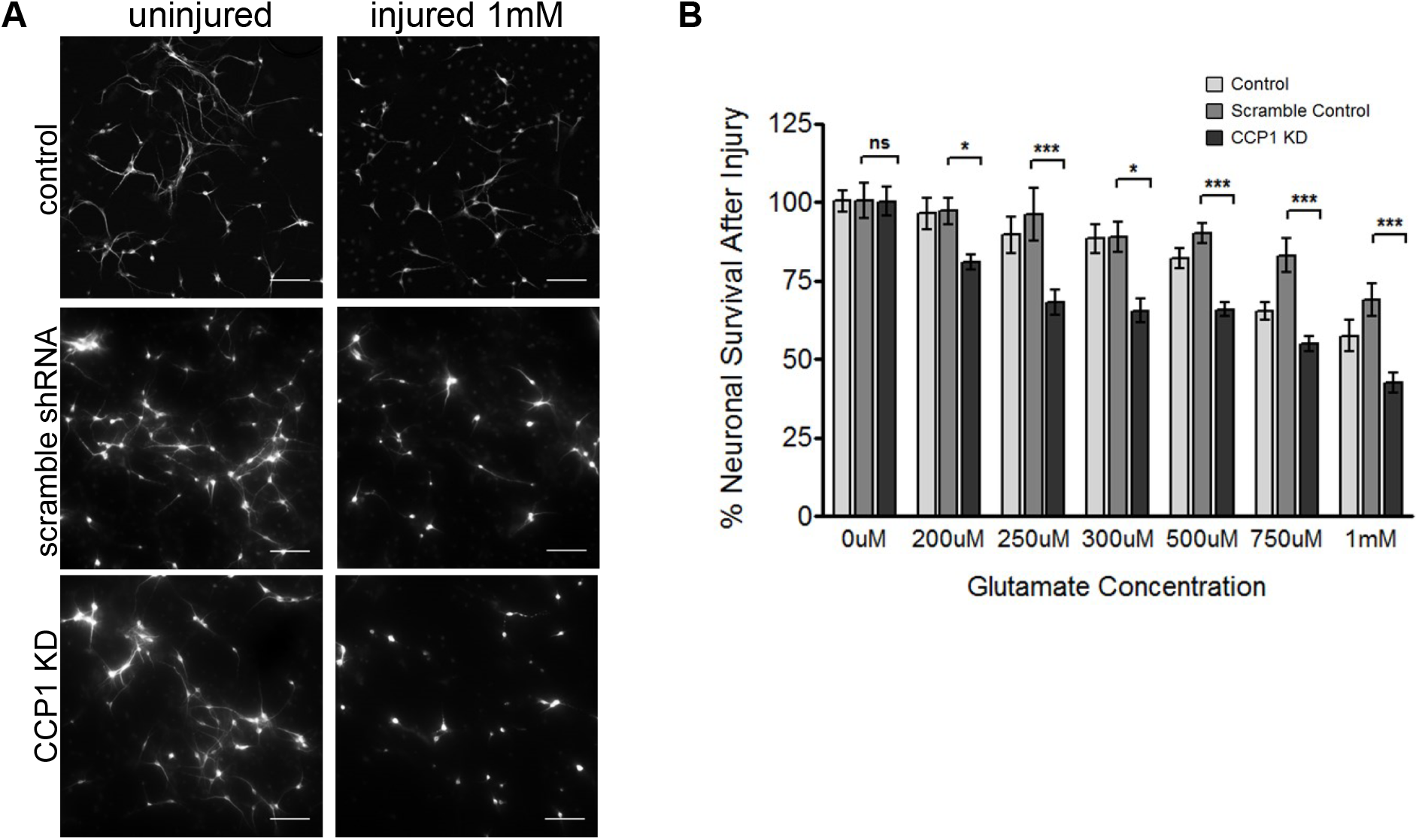
Knockdown of CCP1 decreases neuronal survival after glutamate-induced excitotoxicity *in vitro*. **A.** Representative images of primary spinal cord cultures immunostained for MAP2, before and after injury. Following injury, fewer neurons remained, and their processes were shorter. Scale bars indicate 100μm. **B.** Quantification of percentage of neuronal survival following treatment with concentrations of glutamate ranging from 200μM to 1mM. At every concentration, CCP1 knockdown cultures show significantly reduced neuronal survival compared to scrambled shRNA control, determined by two-way ANOVA followed by Bonferroni post-test analysis. \**p* < 0.05, \*\**p* < 0.01, \*\*\**p* < 0.001. Percentage neuronal survival was not significantly different between control and scramble shRNA control groups.

To confirm that the partial loss of CCP1 is responsible for decreased neuronal survival following glutamate-induced excitotoxicity, we co-infected cultures with two lentiviruses carrying CCP1 shRNA and an shRNA-resistant CCP1 cDNA construct. Western blot analysis confirmed that the levels of CCP1 were rescued to control levels in these cultures (**Fig. 5A-B**). Cultures in which levels of CCP1 were rescued exhibited neuronal survival after excitotoxic challenge comparable to control cultures (**Fig. 5C-D**). We conclude that the presence of CCP1 is needed for neuroprotection from excitotoxicity.

**Fig. 5.**
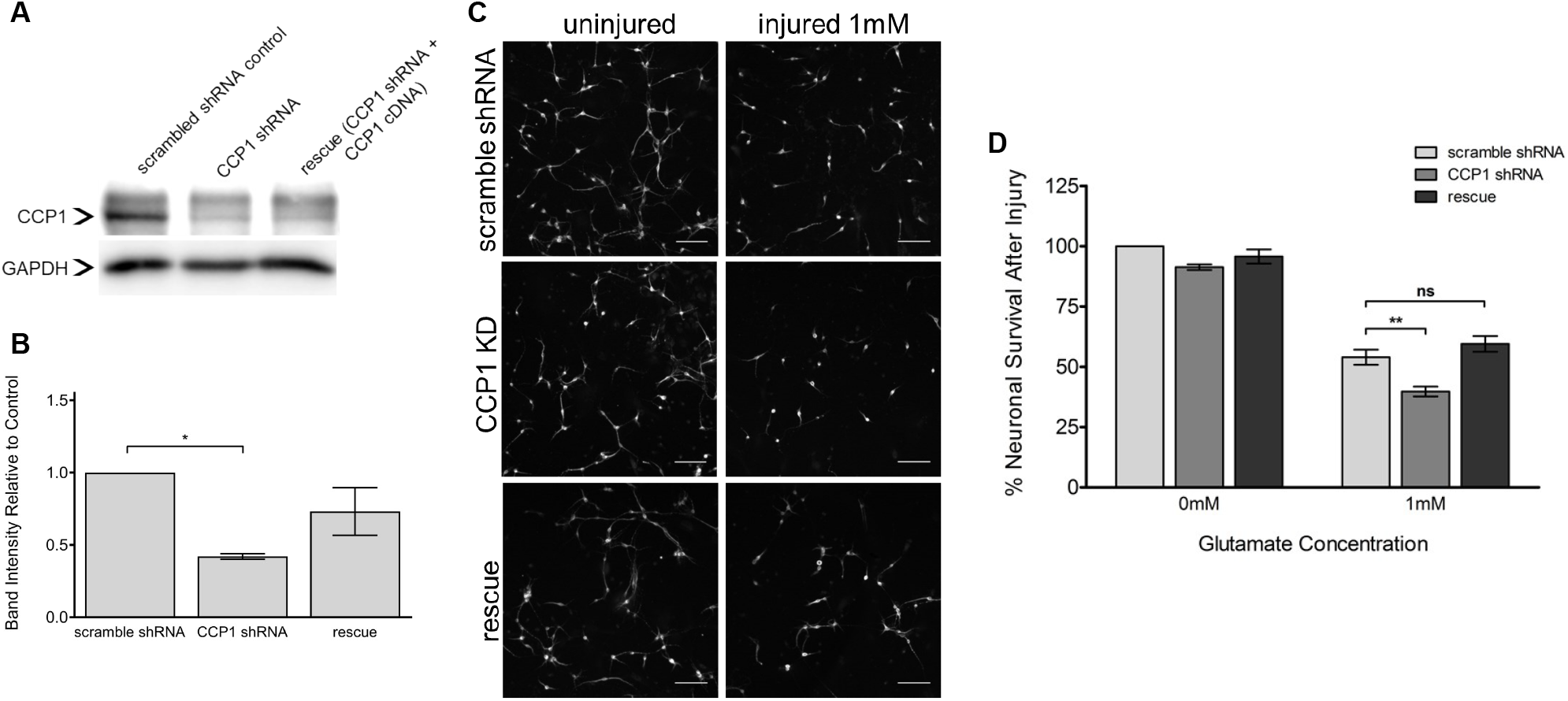
Lentivirus encoding shRNA-resistant CCP1 cDNA rescues decreased neuronal survival following knockdown. **A.** Western blot showing abundance of CCPP1 in scramble, knockdown and rescued cultures. **B.** Quantification Western blot of CCP1 rescue. *p* < 0.05, determined by paired t-test, n=2. **C.** Primary spinal cord cultures immunostained for neuronal marker MAP2 before and after injury. Cultures infected with lentivirus expressing CCP1 following knockdown showed rescue of neuronal death and retraction of processes after injury. Scale bars indicate 100μm. **D.** Quantification of percentage neuronal survival following treatment with 0μM and 1mM glutamate. At both concentrations, neuronal survival of rescue cultures and control cultures are not significantly different, determined by two-way ANOVA followed by Bonferroni post-test analysis; \*\**p* < 0.01.

### CCP1 levels decrease within 24 hours after excitotoxic injury

Because the mechanistic role of CCP1 in neuronal injury response has not been fully characterized, we sought to determine whether levels of CCP1 change after glutamate-induced excitotoxicity. We observed that the levels of CCP1 decrease at 24h after excitotoxic injury to approximately 60% of basal levels by Western blot analysis (**Fig. 6A, B**). In contrast, cultures with CCP1 knockdown did not show an additional decrease in CCP1 levels after glutamate exposure. Because lentiviral infection was more efficient in neurons than in glial cells, our results suggest that the levels of CCP1 protein are tightly regulated in neurons before and after excitotoxic injury to optimize neuronal survival.

**Fig. 6.**
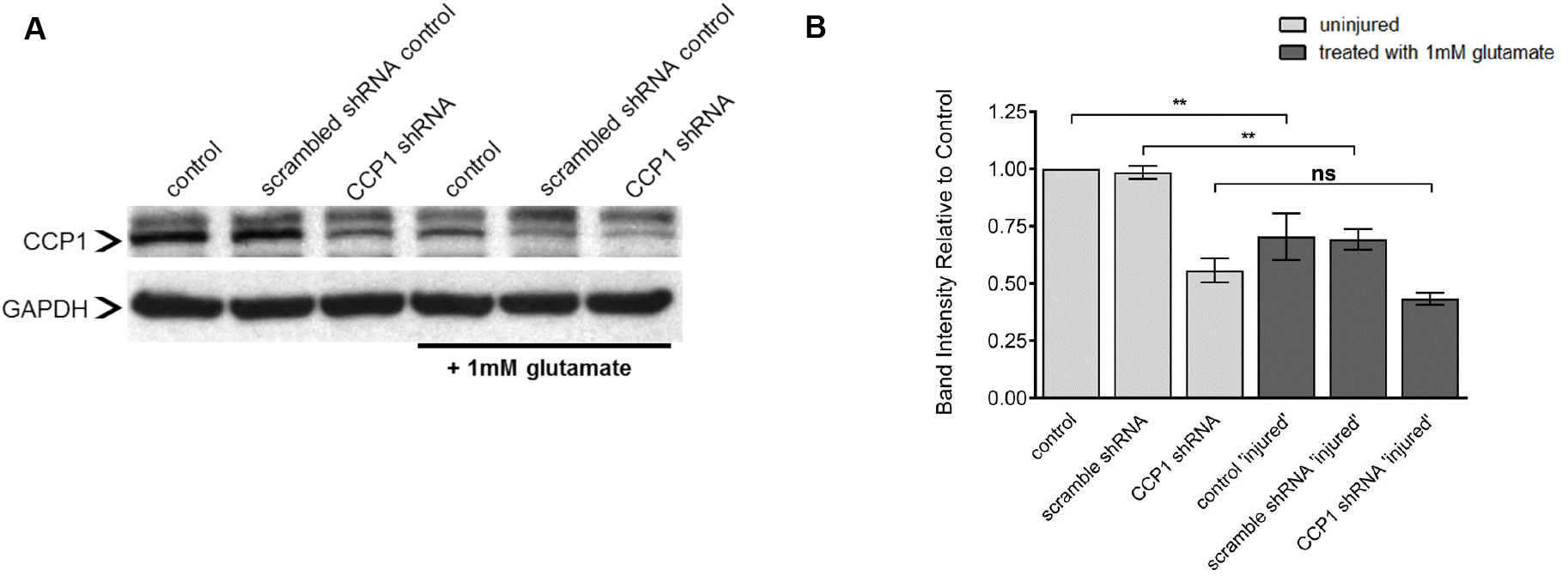
CCP1 is regulated after glutamate-induced excitotoxicity *in vitro.* **A.** Western blot showing changes in CCP1 before and 24 hours after injury. GAPDH was used as a loading control. **B.** Quantification of CCP1 band intensity from panel A. All analyses determined by one-way ANOVA of relative band intensity followed by Tukey’s multiple comparisons test. ** *p* < 0.01, n = 3. See also **Fig. S2.**

### Presence of neuronal cilia does not strongly correlate with survival of excitotoxic injury

Cilia can play a neuroprotective role in the CNS as an antagonist to the cell cycle by inhibiting cell division and preventing apoptosis [40]. To assess if the presence of cilia plays a role in neuronal survival after glutamate-induced excitotoxicity, we compared the fraction of ciliated neurons after treatment with 500μM glutamate to uninjured neurons. We hypothesized that if ciliation is neuroprotective, the frequency of cilia would increase in the neurons remaining after excitotoxic treatment, due to a decreased likelihood of survival of neurons without cilia. We found that in injured cultures, the percentage of neurons that were ciliated in CCP1 knockdown-treated cultures was approximately 69% versus 64% in scramble shRNA-treated controls **(Fig. 7A, B)**. Although the mean percentage was greater in CCP1 knockdown cultures, the difference was not statistically significant (**Fig. 7B**). The length of neuronal cilia was also not significantly different between CCP-1 knockdown and scramble shRNA-treated glutamate-injured cultures **(Fig. 7C)**. Therefore, our data suggest that cilia are not needed for CCP1 to protect sensory spinal cord neurons from excitoxic death.

**Fig. 7.**
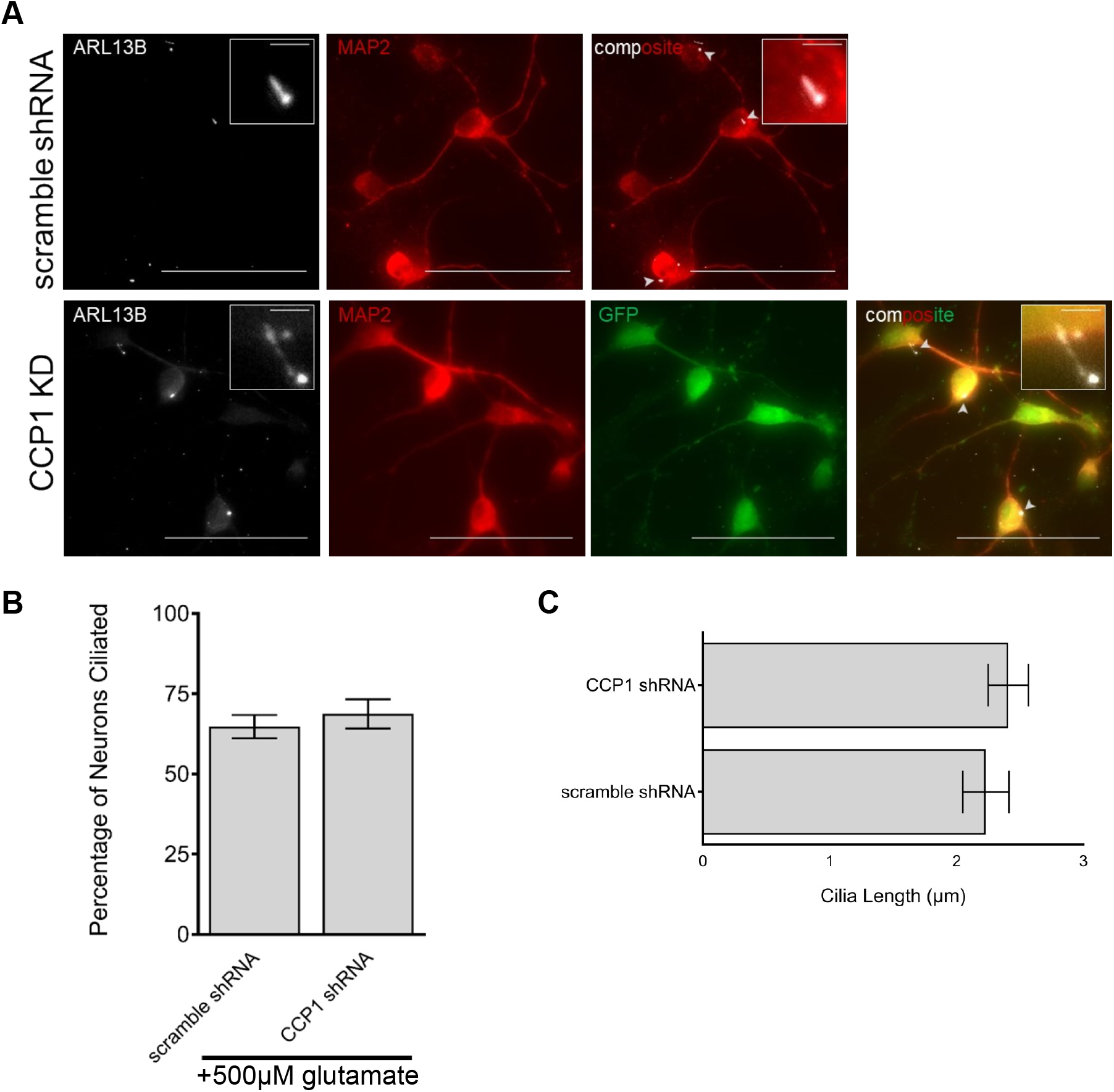
Spinal cord neurons are ciliated post-injury. **A.** Primary spinal cord cultures immunostained for MAP2 and ARL13B, showing ciliated neurons following treatment with 500μM glutamate. Scale bars indicate 50μm and2.5μm (inset). Arrowheads indicate cilia. **B.** Quantification of percentage of neurons ciliated and their respective average lengths in scramble control and CCP1 knockdown cultures. Scramble and knockdown groups were not significantly different, determined by Student’s t-test.

## Discussion

Several lines of evidence support the idea that glutamylation acts a component of the Tubulin Code to regulate the MT cytoskeleton and MT-based motors. Glutamylation is a post-translational modification found on stable MTs [20, 41, 42]. MT glutamylation has been shown to regulate MT-based motor trafficking in neurons of *C. elegans* and mice [13, 14, 21, 22] as well as *in vitro* studies using purified tubulins and motors [43]. Defects in tubulin modifications, MT stability, and neuronal trafficking are linked to neurodegenerative diseases, such as Parkinson’s, Huntington’s, and Alzheimer’s [44–47]. Additionally, in humans, loss of the deglutamylase CCP1 (and resulting hyperglutamylation of MTs) leads to fatal infantile neurodegeneration in the spinal cord, cerebellum, and peripheral nerves [23].

Neuronal expression of CCP1 (also known as NNA1 or AGTBP1) is upregulated after injury of the sciatic nerve, and elevated expression is maintained during target reinnervation [24], suggesting that CCP1 plays a role in neuroregeneration and axonal outgrowth after injury. Therefore, control of MT glutamylation could represent an important survival factor in the context of both neurodegenerative disease and neuronal injury. In order for regeneration after injury to occur, neurons must also survive [48, 49]. In this work, by combining lentivirally-delivered shRNA knockdown combined with a previously established *in vitro* model of the secondary phase of spinal cord injury [26], we show that CCP1 function promotes survival of neurons challenged with excitotoxic levels of glutamate.

We hypothesized that CCP1 might function in cilia to promote survival of neurons exposed to excitotoxic injury. Cilia can play a neuroprotective role in the rodent CNS [40] and are important for reception of cellular signaling that is essential for development of the mammalian nervous system [9, 11, 50]. Motile cilia on ependymal cells, such as those that line the central canal, are essential for spinal cord morphogenesis [51]. Cerebrospinal fluid-contacting neurons, which also have a motile cilium, extend an apical projection into the central canal and are proposed to relay cerebrospinal fluid flow and pH information spinal circuits for normal development and function of spinal cord nerves [32, 35, 52]. Spinal cord injury can cause degeneration of motile ependymal cilia or ependymal cells, which create cerebrospinal fluid flow in the central canal and neurons [53]. This could result in toxic buildup of CNS waste, possibly preventing regrowth and exacerbating the chronic degeneration of spinal tissues after SCI [53]. CCP1 homologs have been found to regulate the integrity and structure of MTs in cilia, the function of ciliary motors [13, 14, 27, 54, 55], and ciliary length [31].

To our knowledge, no previous reports had found that sensory neurons or interneurons of the spinal cord were ciliated. Spinal cord motorneurons have been shown to be ciliated *in vitro* [37]. We used immunofluorescence to detect the ciliary protein ARL13B and found that neurons in embryonic spinal cord cultures are ciliated. However, under the culture conditions we used, spinal cord cultures lack motorneurons (**Fig. S1A**, [39]). Therefore, our data provide the first evidence (to our knowledge) that spinal cord sensory neurons are ciliated.

Our data suggest that expression of CCP1 in the context of the spinal cord might not strongly regulate ciliation. This was surprising, as we had previously demonstrated that the lack of a CCP1 homolog in nematodes causes the degeneration of neuronal cilia [13]. An siRNA screen in immortalized human retinal pigmented epithelial cells had also found that CCP1 can positively regulate cilia length [31]. Our result may be explained by the fact that the Tubulin Code, and glutamylation in particular, can result in cell-specific effects [13, 14], likely due to differences in expression of genes that read or interpret the Tubulin Code modifications. Because we did not detect a clear relationship between the presence of cilia and neuronal survival, we suggest that CCP1 may promote neuronal survival mainly by deglutamylating axonal and dendritic, rather than ciliary, MTs. However, using our *in vitro* system, we cannot address how CCP1 might function in cilia of ependymal cells, cerebrospinal fluid-contacting neurons, or motorneurons in responses to nerve injury.

Because CCP1 deglutamylates MTs [16], we hypothesize that regulation of CCP1 expression and function is necessary to appropriately reorganize MT dynamics following neuronal injury to facilitate axonal regeneration and recovery. This notion is supported by the previous finding that CCP1 homologs in the invertebrate *C. elegans* are necessary for normal neuronal outgrowth following axotomy of touch receptor neurons [25]. When CNS axons fail to regenerate, MTs are disorganized after injury, whereas regenerating PNS axons form organized MT networks in their growth cones [56]. Additionally, MT defects are proposed to underlie neurodegeneration after traumatic brain injury [57].

Our results suggest that CCP1 function is needed before and during excitotoxic injury to improve neuronal survival. We found that CCP1 expression significantly decreased following glutamate-induced excitotoxicity in spinal cord neurons. When CCP1 was knocked down before injury, excitotoxicity did not further reduce CCP1 expression. Our observations differ from a previous study [24], which shows upregulation of CCP1 following transection or nerve crush injury of the sciatic nerve in the PNS. There are at least two possible explanations for the observed difference in CCP1 levels. Acute injury that severs or damages the axonal cytoskeleton might increase CCP1 expression, while chronic (excitotoxic) injury might decrease CCP1 expression. Alternatively, injury of PNS neurons might upregulate CCP1 expression, while injury of the CNS might downregulate of CCP1 expression. In this case, differences in the function or structure of the MT cytoskeleton, mediated at least in part by CCP1, might explain the poor regenerative potential in the CNS and robust regeneration in the PNS.

Our results support the notion that therapeutic targeting of intrinsic neuronal molecules that regulate MTs, such as CCP1, along with drugs targeting extrinsic inhibitors of regrowth, might offer an effective combination for improving neuroregeneration after spinal cord injury in the future.

## Acknowledgments

We thank Anton Olmechenko for help with spinal cord culture techniques and members of the Barr and Firestein labs for helpful discussions. This work was funded by NJCSCR grant CSCR15IRG014 (to R.O.), NIH grants DK059418 and DK074746 (to M.M.B.), and NJCSCR grants CSCR14IRG005 and CSCR17ERG005 (to B.L.F).

---

**Fig. S1.**
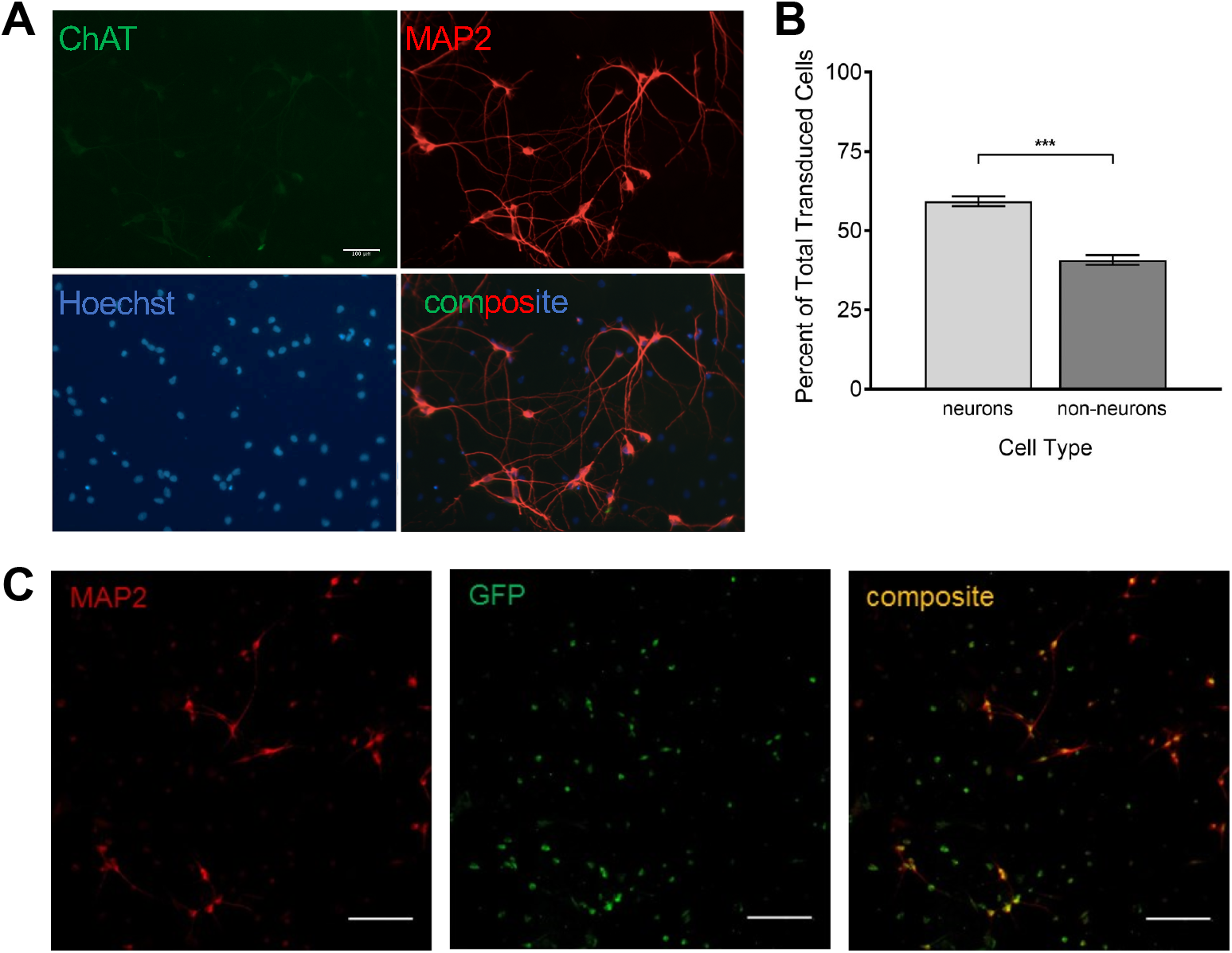
shRNA-mediated knock-down of CCP1 affected neurons. **A.** Representative images of MAP2 and ChAT immunostaining of mixed spinal cord cultures. Lack of ChAT immunostaining indicates that the neuronal population is not motor but most likely sensory. **B.** Quantification of percentage of transduction by cell type. **C.** Representative images of primary spinal cord cultures infected with a lentivirus carrying scrambled shRNA or CCP1 shRNA. Neurons were immunostained for MAP2, and infected neurons express GFP as a fluorescent transduction marker. Scale bars indicate 100μm.

**Fig. S2.**
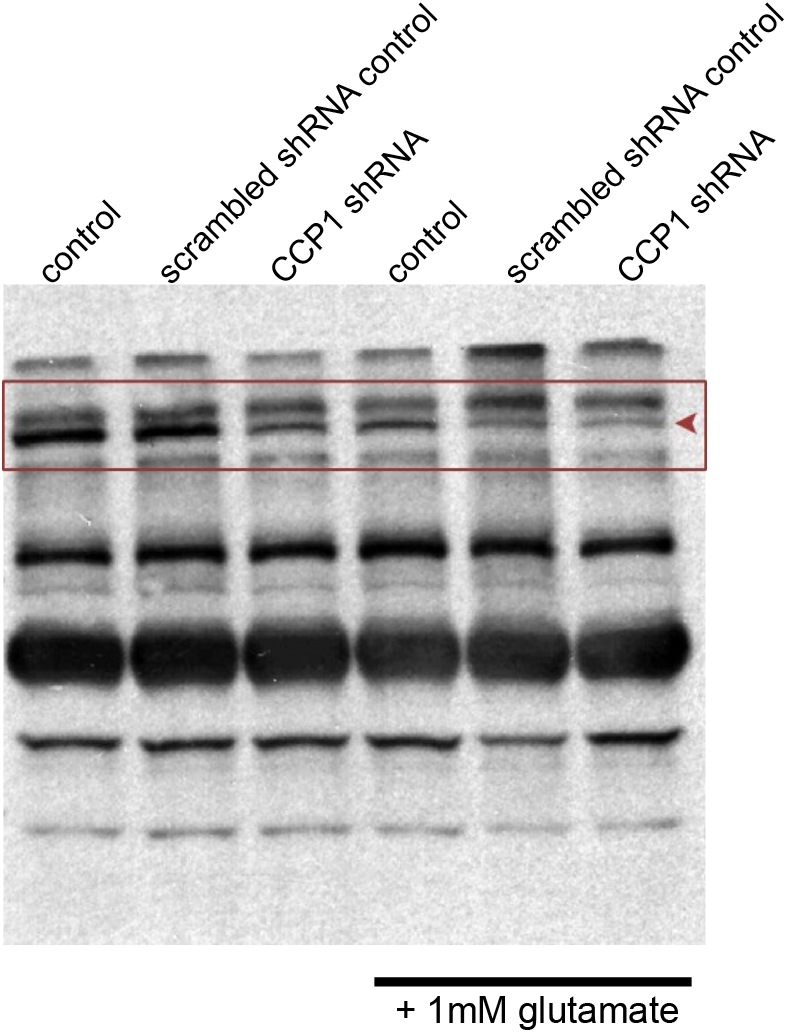
Results of binding of CCP1 antibody in Western blot assays. Arrow indicates CCP1 bands at 137kDa, identifiable by size and pattern across control and CCP1 knockdown groups. Other bands did not change across groups and were considered non-specific cross-reacting proteins.

## Materials and Methods

All animal experiments were conducted in accordance with the National Institutes of Health (NIH) Guide for the Care and Use of Laboratory animals (NIH Publication No. 8023, revised 1978).

### Spinal Cord Cultures

Our method was similar to that used by Du *et al.,* 2007 [26]. Briefly, spinal cords were dissected from Sprague-Dawley rat embryos at gestational day 15 (E15) by removing meninges and attached dorsal root ganglia. Cords were then gently triturated to dissociate a mixture of neurons, astrocytes, and microglia from the tissue. Cells were plated at a density of approximately 523 cells/mm^2^, or 100,000 cells per well, in a 24 well plate containing coverslips coated with 0.01% solution of poly-d-lysine (Sigma) dissolved in 0.1M borate buffer (sodium tetraborate, boric acid) for cell adhesion. Mixed cultures were grown in Dulbecco’s Modified Eagle Medium (DMEM; Gibco) supplemented with 10% Horse Serum (Gibco) for 7 days before glutamate treatment.

### Lentiviral production & infection

Lentiviruses were produced as previously described [58]. In summary, HEK293T cells, plated at a density of 6.5*10^6^ cells/T75 flask were grown for one day and then transfected on DIV2 (days *in vitro* 2) with lentiviral plasmids (VectorBuilder) carrying scrambled shRNA (target sequence: CCTAAGGTTAAGTCGCCCTCG; Vector ID: VB170329-1128paq), CCP1 knockdown shRNA (target sequence: CCACTTCCAGTTGCCAATTAT; Vector ID: VB171011-1192pab) or CCP1 cDNA (RGD ID: 1306307; AGTPBP1 also known as CCP1) and GFP or RFP fluorescent markers. VSV and PAX2 packaging vectors and Lipofectamine 2000 (Invitrogen) were used according to the manufacturer’s protocol to mediate transfection. After three days, the media were collected and centrifuged at 1500 × *g* for 5 minutes to pellet dead cells and debris. The supernatant was then collected and incubated with PEG-it™ (System Biosciences) in a 5X solution at 4C for 2 days to precipitate the viruses. On day 7, the solution was centrifuged at 1500 × *g* for 30 minutes at 4C to pellet the viruses. The supernatant was removed and discarded, and the virus pellet was resuspended in 150μl of sterile 1X PBS and frozen (at −80°C) in 5μl aliquots until use.

All spinal cord culture infections were performed on day *in vitro* 2 (DIV2) at a 1:5000 dilution by replacing one-fourth of the medium in the well with a 1:1250 dilution of virus in fresh DMEM + 10% Horse Serum.

### Glutamate Treatment

Glutamate-induced excitotoxicity was performed as previously described [26]. Glutamate (L-glutamic acid, Sigma Life Sciences) dilutions were made from a 1mM stock solution dissolved in DMEM + 10% Horse Serum warmed to 37C. During treatment on DIV7, medium was collected from each well and replaced with varying concentrations of glutamate-containing medium as described in results for 1 hour at 37°C and 5% CO_2_. Conditioned medium (collected before treatment) was combined with an equal volume of fresh medium (1:1 solution) and used as recovery medium following glutamate treatment. Cells were allowed to recover for 24 hours at 37°C and 5% CO_2_ before fixation.

### Immunocytochemistry

Cells were fixed by incubation in 4% paraformaldehyde in 1X phosphate-buffered saline (PBS) for 15 minutes. Following fixation, cells were washed three times with 1X PBS and incubated in a blocking solution (2% Normal Goat Serum, 0.1% Triton-X 100, 0.02% NaN_3_ diluted in 1X PBS) for 1 hour. Cells were incubated in primary antibody solution (1:500) overnight (approximately 18 hours), washed three times in 1X PBS, and incubated with appropriate secondary antibody solutions (1:1000) for 1 hour the next day, followed by another three washes in 1X PBS. Coverslips were then removed and mounted on glass slides for imaging.

Anti-MAP2 antibody (microtubule associated protein 2; Thermofisher) was used to identify neurons. Anti-GFAP (glial fibrillary acidic protein; Millipore, rabbit) was used to label astrocytes, and anti-IBA1 (ionized calcium binding adaptor molecule 1; Proteintech, mouse) was used to label microglia. Cilia were immunolabeled using a monoclonal anti-ARL13B antibody (Proteintech, mouse). Anti-ChAT (Choline Acetyltransferase; Millipore Sigma, goat) antibody was used to identify motorneurons.

AlexaFluor-conjugated secondary antibodies (488, 568, 647) were used to visualize cell-specific markers. Nuclei were labelled using NucBlue Live Ready Probes Reagent (Thermofisher), which was excited at 350nm.

### Imaging and Microscopy

Image Z-stacks were acquired with Metamorph software (www.moleculardevices.com) using a Zeiss Axioplan2 microscope with 10×, 63× (NA 1.4), and 100× (NA 1.4) oil-immersion objectives, equipped with a Hamamatsu C11440-42U ORCA-Flash4.0 LT Digital CMOS camera (www.hamamatsu.com). Images were uploaded into FIJI/Image J 2.0 to create optical Z-stack projections, add scale bars, and adjust brightness/contrast. The cell counter plugin was used to count MAP2+ and GFP+ cells. Images were then exported as PNG files for assembly into figures in Adobe Illustrator CS.

### Scoring Neuronal Survival

To assess neuronal survival following glutamate treatment, neurons identified by MAP2 immunostaining were counted from five 1.4μm × 1.4μm regions imaged from each coverslip using the 100x objective and analyzed as ratios of neurons remaining following glutamate treatment/ neurons present in the absence of a glutamate treatment.

### Scoring Neuronal Ciliation and Cilia Length

Neurons were identified using anti-MAP2 immunofluorescence, and viral infection with shRNA vectors (scrambled or anti-CCP1) was scored by expression of GFP. For ciliation, the presence or absence of a cilium immunolabeled by ARL13B on the cell body of each neuron in 5 randomly chosen areas was scored and analyzed from Z-stacks of images taken using the 100x objective on the Zeiss Axioplan2 microscope. Z-stacks were uploaded into FIJI/ImageJ, Z-projected, and neuronal cilia were counted using the Cell Counter plugin.

For cilia length, only neurons with ‘horizontally-projecting’ cilia (the entire length of the cilium was visible in a single focal plane of a Z-stack) were scored from images taken with the 100x objective on the Zeiss Axioplan2 microscope. Z-stacks were uploaded into FIJI/ImageJ, Z-projected, and measured for pixel length using the tracing tool. Pixels were converted to microns (at 100x, 0.6566um = 1 pixel) before graphing & statistical analysis.

### Western Blot Analysis

For Western blot assays, spinal cord cultures were grown at 1 million cells per well in a 6 well plate and infected with viruses and/or treated with glutamate at similar concentrations as described above. On DIV8, cells were homogenized in RIPA buffer (50mM Tris-HCl, pH 7.4, 150mM NaCl, 0.5% deoxycholate, 1% NP-40, 0.1% sodium dodecyl sulfate (SDS), 1mM ethylenediaminetetraacetic acid (EDTA), pH 7.4) by scraping the cells into the buffer and sonication. Protein concentrations were determined using the Pierce BCA protein assay kit (Thermo Scientific) according to the manufacturer’s protocol. Proteins (30μg/lane) were resolved on a 10% SDS polyacrylamide gel and transferred to a polyvinylidene difluoride (PVDF) membrane. Membranes were blocked with 5% bovine serum albumin (BSA) in 1X TBST (20mM Tris pH 7.5, 150mMNaCl, 0.1% Tween-20) for 1 hour. Blots were incubated in primary antibody solutions (1:1000) overnight and then washed and incubated in appropriate horseradish peroxidase-conjugated secondary antibody solutions (1:5000, Rockland) for 1 hour the next day. Blots were probed with the following antibodies: anti-CCP1 (Proteintech, rabbit) and anti-glyceraldehyde 3-phosphate dehydrogenase (GAPDH; EMD Millipore, mouse), which served as a loading control. Bands were visualized using the LI-COR Biosciences digital imaging system and pixel intensity was analyzed using ImageJ (NIH). CCP1 band intensities were normalized to corresponding GAPDH band intensities.

### Experimental Design and Statistical Analyses

All statistical analyses were performed using a combination of GraphPad Prism (Version 5.01, GraphPad Software, La Jolla, CA, USA) and Microsoft Excel (Version 14.0.7106, 32-Bit, Microsoft Corporation, Seattle, WA, USA).

## References

1. Varidaki, A., Y. Hong, and E.T. Coffey, Repositioning Microtubule Stabilizing Drugs for Brain Disorders. Front Cell Neurosci, 2018. 12: p. 226.

2. Witte, H. and F. Bradke, The role of the cytoskeleton during neuronal polarization. Curr Opin Neurobiol, 2008. 18(5): p. 479–87.

3. Pellegrini, L., et al., Back to the tubule: microtubule dynamics in Parkinson’s disease. Cell Mol Life Sci, 2017. 74(3): p. 409–434.

4. Gadadhar, S., et al., The tubulin code at a glance. J Cell Sci, 2017. 130(8): p. 1347–1353.

5. Verhey, K.J. and J. Gaertig, The tubulin code. Cell Cycle, 2007. 6(17): p. 2152–60.

6. Edde, B., et al., Posttranslational glutamylation of alpha-tubulin. Science, 1990. 247(4938): p. 83–5.

7. Bre, M.H., et al., Glutamylated tubulin probed in ciliates with the monoclonal antibody GT335. Cell Motil Cytoskeleton, 1994. 27(4): p. 337–49.

8. Werner, S., A. Pimenta-Marques, and M. Bettencourt-Dias, Maintaining centrosomes and cilia. J Cell Sci, 2017. 130(22): p. 3789–3800.

9. Youn, Y.H. and Y.G. Han, Primary Cilia in Brain Development and Diseases. Am J Pathol, 2018. 188(1): p. 11–22.

10. Lee, J.E. and J.G. Gleeson, Cilia in the nervous system: linking cilia function and neurodevelopmental disorders. Curr Opin Neurol, 2011. 24(2): p. 98–105.

11. Bay, S.N. and T. Caspary, What are those cilia doing in the neural tube? Cilia, 2012. 1(1): p. 19.

12. Ikegami, K. and M. Setou, Unique post-translational modifications in specialized microtubule architecture. Cell Struct Funct, 2010. 35(1): p. 15–22.

13. O’Hagan, R., et al., The tubulin deglutamylase CCPP-1 regulates the function and stability of sensory cilia in C. elegans. Current biology : CB, 2011. 21(20): p. 1685–94.

14. O’Hagan, R., et al., Glutamylation Regulates Transport, Specializes Function, and Sculpts the Structure of Cilia. Curr Biol, 2017. 27(22): p. 3430–3441 e6.

15. Rodriguez de la Vega, M., et al., Nna1-like proteins are active metallocarboxypeptidases of a new and diverse M14 subfamily. Faseb J, 2007. 21(3): p. 851–65.

16. Rogowski, K., et al., A family of protein-deglutamylating enzymes associated with neurodegeneration. Cell, 2010. 143(4): p. 564–78.

17. Power, K.A., JS; Gu, A; Walsh, JD; Bellotti, S; Morash, M; Zhang, W; Ross, N; Golden, A; Smith, HE; Barr, MM; O’Hagan, R Mutation of NEKL-4/NEK10 and TTLL genes opposes loss of the CCPP-1 deglutamylase and prevents neuronal ciliary degeneration. biorxiv.org, 2020. DOI: https://doi.org/10.1101/2020.05.21.108449.

18. Fernandez-Gonzalez, A., et al., Purkinje cell degeneration (pcd) phenotypes caused by mutations in the axotomy-induced gene, Nna1. Science, 2002. 295(5561): p. 1904–6.

19. Mitchison, H.M. and E.M. Valente, Motile and non-motile cilia in human pathology: from function to phenotypes. J Pathol, 2017. 241(2): p. 294–309.

20. Fukushima, N., et al., Post-translational modifications of tubulin in the nervous system. J Neurochem, 2009. 109(3): p. 683–93.

21. Ikegami, K., et al., Loss of alpha-tubulin polyglutamylation in ROSA22 mice is associated with abnormal targeting of KIF1A and modulated synaptic function. Proc Natl Acad Sci U S A, 2007. 104(9): p. 3213–8.

22. Magiera, M.M., et al., Excessive tubulin polyglutamylation causes neurodegeneration and perturbs neuronal transport. EMBO J, 2018. 37(23).

23. Shashi, V., et al., Loss of tubulin deglutamylase CCP1 causes infantile-onset neurodegeneration. EMBO J, 2018. 37(23).

24. Harris, A., et al., Regenerating motor neurons express Nna1, a novel ATP/GTP-binding protein related to zinc carboxypeptidases. Mol Cell Neurosci, 2000. 16(5): p. 578–96.

25. Ghosh-Roy, A., et al., Kinesin-13 and tubulin posttranslational modifications regulate microtubule growth in axon regeneration. Developmental cell, 2012. 23(4): p. 716–28.

26. Du, Y., et al., Astroglia-mediated effects of uric acid to protect spinal cord neurons from glutamate toxicity. Glia, 2007. 55(5): p. 463–72.

27. Kubo, T., et al., Reduced tubulin polyglutamylation suppresses flagellar shortness in Chlamydomonas. Mol Biol Cell, 2015. 26(15): p. 2810–22.

28. Campbell, P.K., et al., Mutation of a novel gene results in abnormal development of spermatid flagella, loss of intermale aggression and reduced body fat in mice. Genetics, 2002. 162(1): p. 307–20.

29. Branham, K., et al., Establishing the involvement of the novel gene AGBL5 in retinitis pigmentosa by whole genome sequencing. Physiol Genomics, 2016. 48(12): p. 922–927.

30. Chakrabarti, L., et al., The zinc-binding domain of Nna1 is required to prevent retinal photoreceptor loss and cerebellar ataxia in Purkinje cell degeneration (pcd) mice. Vision research, 2008. 48(19): p. 1999–2005.

31. Kim, J., et al., Functional genomic screen for modulators of ciliogenesis and cilium length. Nature, 2010. 464(7291): p. 1048–51.

32. Bohm, U.L., et al., CSF-contacting neurons regulate locomotion by relaying mechanical stimuli to spinal circuits. Nat Commun, 2016. 7: p. 10866.

33. Djenoune, L., et al., Investigation of spinal cerebrospinal fluid-contacting neurons expressing PKD2L1: evidence for a conserved system from fish to primates. Front Neuroanat, 2014. 8: p. 26.

34. Orts-Del’Immagine, A., et al., Morphology, distribution and phenotype of polycystin kidney disease 2-like 1-positive cerebrospinal fluid contacting neurons in the brainstem of adult mice. PLoS One, 2014. 9(2): p. e87748.

35. Sternberg, J.R., et al., Pkd2l1 is required for mechanoception in cerebrospinal fluid-contacting neurons and maintenance of spine curvature. Nat Commun, 2018. 9(1): p. 3804.

36. Meletis, K., et al., Spinal cord injury reveals multilineage differentiation of ependymal cells. PLoS Biol, 2008. 6(7): p. e182.

37. Ma, X., R. Peterson, and J. Turnbull, Adenylyl cyclase type 3, a marker of primary cilia, is reduced in primary cell culture and in lumbar spinal cord in situ in G93A SOD1 mice. BMC Neurosci, 2011. 12: p. 71.

38. Higginbotham, H., et al., Arl13b in primary cilia regulates the migration and placement of interneurons in the developing cerebral cortex. Dev Cell, 2012. 23(5): p. 925–38.

39. Kushima, Y. and H. Hatanaka, Interleukin-6 and leukemia inhibitory factor promote the survival of acetylcholinesterase-positive neurons in culture from embryonic rat spinal cord. Neurosci Lett, 1992. 143(1-2): p. 110–4.

40. Choi, B.K.A., et al., Stabilization of primary cilia reduces abortive cell cycle re-entry to protect injured adult CNS neurons from apoptosis. PLoS One, 2019. 14(8): p. e0220056.

41. Janke, C. and J.C. Bulinski, Post-translational regulation of the microtubule cytoskeleton: mechanisms and functions. Nature reviews. Molecular cell biology, 2011. 12(12): p. 773–86.

42. Wloga, D., E. Joachimiak, and H. Fabczak, Tubulin Post-Translational Modifications and Microtubule Dynamics. Int J Mol Sci, 2017. 18(10).

43. Sirajuddin, M., L.M. Rice, and R.D. Vale, Regulation of microtubule motors by tubulin isotypes and post-translational modifications. Nat Cell Biol, 2014. 16(4): p. 335–44.

44. Vu, H.T., et al., Increase in alpha-tubulin modifications in the neuronal processes of hippocampal neurons in both kainic acid-induced epileptic seizure and Alzheimer’s disease. Sci Rep, 2017. 7: p. 40205.

45. Brady, S.T. and G.A. Morfini, Regulation of motor proteins, axonal transport deficits and adult-onset neurodegenerative diseases. Neurobiol Dis, 2017. 105: p. 273–282.

46. Matamoros, A.J. and P.W. Baas, Microtubules in health and degenerative disease of the nervous system. Brain Res Bull, 2016. 126(Pt 3): p. 217–225.

47. Baas, P.W., et al., Stability properties of neuronal microtubules. Cytoskeleton (Hoboken), 2016. 73(9): p. 442–60.

48. Dusart, I., et al., Cell death and axon regeneration of Purkinje cells after axotomy: challenges of classical hypotheses of axon regeneration. Brain Res Brain Res Rev, 2005. 49(2): p. 300–16.

49. Hollis, E.R., 2nd, et al., IGF-I gene delivery promotes corticospinal neuronal survival but not regeneration after adult CNS injury. Exp Neurol, 2009. 215(1): p. 53–9.

50. Louvi, A. and E.A. Grove, Cilia in the CNS: the quiet organelle claims center stage. Neuron, 2011. 69(6): p. 1046–60.

51. Grimes, D.T., et al., Zebrafish models of idiopathic scoliosis link cerebrospinal fluid flow defects to spine curvature. Science, 2016. 352(6291): p. 1341–4.

52. Djenoune, L., et al., The dual developmental origin of spinal cerebrospinal fluid-contacting neurons gives rise to distinct functional subtypes. Sci Rep, 2017. 7(1): p. 719.

53. Radojicic, M., G. Nistor, and H.S. Keirstead, Ascending central canal dilation and progressive ependymal disruption in a contusion model of rodent chronic spinal cord injury. BMC neurology, 2007. 7: p. 30.

54. Suryavanshi, S., et al., Tubulin glutamylation regulates ciliary motility by altering inner dynein arm activity. Current biology : CB, 2010. 20(5): p. 435–40.

55. Hong, S.R., et al., Spatiotemporal manipulation of ciliary glutamylation reveals its roles in intraciliary trafficking and Hedgehog signaling. Nat Commun, 2018. 9(1): p. 1732.

56. Erturk, A., et al., Disorganized microtubules underlie the formation of retraction bulbs and the failure of axonal regeneration. J Neurosci, 2007. 27(34): p. 9169–80.

57. Tang-Schomer, M.D., et al., Mechanical breaking of microtubules in axons during dynamic stretch injury underlies delayed elasticity, microtubule disassembly, and axon degeneration. FASEB J, 2010. 24(5): p. 1401–10.

58. Patel, M.V., et al., A Role for Postsynaptic Density 95 and Its Binding Partners in Models of Traumatic Brain Injury. J Neurotrauma, 2019. 36(13): p. 2129–2138.

